# Description of two novel *Corynebacterium* species isolated from human nasal passages and skin

**DOI:** 10.1101/2024.11.21.624533

**Authors:** Elena B. Popowitch, Tommy H. Tran, Isabel Fernandez Escapa, Eeshta Bhatt, Alp K. Sozat, Nashwa Ahmed, Clayton Deming, Ari Q. Roberts, NISC Comparative Sequencing Program, Julia A. Segre, Heidi H. Kong, Sean Conlan, Katherine P. Lemon, Matthew S. Kelly

## Abstract

Strains of two novel *Corynebacterium* species were cultured from samples of human nostrils and skin collected in the United States and Botswana. These strains demonstrated growth on Columbia Colistin-Nalidixic Acid agar with 5% sheep blood and in liquid media (brain heart infusion and tryptic soy broth) supplemented with Tween 80, a source of the fatty acid oleic acid. Cells were Gram-positive, non-spore-forming, non-motile bacilli that showed catalase but not oxidase activity. Major fatty acids in both of these species were 18:1 ω9c (oleic acid), 16:0 (palmitic acid), and 18:0 (stearic acid). Analysis of the 16S ribosomal RNA gene sequences identified these strains as belonging to the genus *Corynebacterium* (family Corynebacteriaceae). Whole-genome sequencing revealed that these strains formed distinct branches on a phylogenomic tree, with *C. tuberculostearicum* being the closest relative but with average nucleotide identities of < 95% relative to all previously described species. These results indicate that these strains represent novel species of *Corynebacterium*, for which we propose the names *Corynebacterium hallux* sp. nov., with the type strain CTNIH22^T^ (=ATCC TSD-435^T^=DSM 117774^T^), and *Corynebacterium nasorum* sp. nov., with the type strain KPL3804^T^ (=ATCC TSD-439^T^=DSM 117767^T^). We also describe the characteristics of two strains isolated from human nasal passages that are members of the recently named species *Corynebacterium yonathiae*.

**Repositories:** The sequencing files supporting the conclusions of this study are available in the Sequence Read Archive (PRJNA804245, PRJNA854648, PRJNA842433). The partial 16S ribosomal RNA gene sequences from PCR amplification and Sanger sequencing are available in GenBank for *Corynebacterium hallux* sp. nov. CTNIH22^T^ (accession number: PQ252679) and *Corynebacterium nasorum* sp. nov. KPL3804^T^ (accession number: PQ149068). The annotated genomic sequences for the strains characterized in this study have been deposited in GenBank with the following accession numbers: *C. hallux* sp. nov. CTNIH22^T^ (GCF_032821755.1), *C. nasorum* sp. nov. KPL3804^T^ (GCF_037908315.1), *C. nasorum* sp. nov. MSK185 (GCF_030229765.1), *C. yonathiae* KPL2619 (GCF_037908465.1), and *C. yonathiae* MSK136 (GCF_022288805.2).

## Introduction

The genus *Corynebacterium* belongs to the family *Corynebacteriaceae* and includes more than 150 validly published species. Most *Corynebacterium* species are only rarely associated with disease among humans and animals. Common pathogenic members include *C. diphtheriae*, the causative agent of the human disease diphtheria, and *C. pseudotuberculosis* and *C. ulcerans*, which are frequent causes of zoonotic infections. Historically, species of the genus *Corynebacterium* have been differentiated based on their host, ecological niche, biochemical characteristics, spectrometric analyses [e.g., matrix-assisted laser desorption/ionization time-of-flight mass spectrometry (MALDI-TOF MS)], or sequencing of specific genetic loci [1–3]. With regard to the latter, phylogenies based only on the 16S ribosomal RNA (rRNA) gene often have poor support within this genus, with improved results for phylogenies based on full or partial *rpoB* gene sequences [4]. Multi-locus sequence typing of housekeeping genes is frequently used to characterize common pathogens such as *C. diphtheriae* [5–8]. As a result of decreasing sequencing costs and improved tools for genomic analysis, whole-genome sequencing is increasingly being performed for taxonomic classification of *Corynebacterium* strains.

*C. tuberculostearicum* is a lipid-requiring species that is a common inhabitant of human skin. This species was first described in 1984 when Brown and colleagues identified 16 strains that had biochemical properties that distinguished them from previously described members of the *Corynebacterium* genus [9]. They called this novel species *C. tuberculostearicum* because strains were noted to contain tuberculostearic acid on fatty acid profiling [9]. In 2004, Feurer and colleagues emended the description of *C. tuberculostearicum* and formally proposed it as a new species [10]. In the present study, we identified several *Corynebacterium* strains from human nasal and skin samples that are most closely related to *C. tuberculostearicum*, but that represent distinct species based on their biological properties, chemical structures, and genomic sequences. We propose classification of these strains into novel species *Corynebacterium hallux* sp. nov. (“hallux” referring to the innermost toe) and *C. nasorum* sp. nov. (“nasorum” referring to “of noses”) to reflect the ecological niches from which these strains were isolated. Finally, we provide a detailed characterization of two additional strains isolated from human nasal passages that are members of the recently described species *C. yonathiae* [11].

### Isolation and Ecology

The new *Corynebacterium* strains described in this study are as follows by species: *C. hallux* sp. nov. (CTNIH22^T^=ATCC TSD-435^T^=DSM 117774^T^), *C. nasorum* sp. nov. (KPL3804^T^=ATCC TSD-439^T^=DSM 117767^T^, MSK185), and *C. yonathiae* (KPL2619, MSK136). *C. hallux* sp. nov. strain CTNIH22^T^ was isolated from the toe web of a healthy adult volunteer in 2019. This sample was collected using an ESwab (Copan, Murrieta, CA), placed in liquid Amies transport medium, and grown on brain heart infusion (BHI) agar with 10% Tween 80 at 35°C under aerobic conditions [12]. The KPL strains of *C. nasorum* and *C. yonathiae* were isolated from nasal samples collected from a child and an adult participating in scientific outreach events in Massachusetts in 2017 and 2018, respectively [13]. These samples were inoculated onto plates containing either BBL Columbia Colistin-Nalidixic Acid (CNA) agar with 5% sheep blood or BHI agar with 1% Tween 80 and 25 µg/mL fosfomycin. Cultures were incubated aerobically for 48 hours at 37°C in either atmospheric conditions or 5% carbon dioxide (CO_2_). For suspected *Corynebacterium*, Sanger sequencing was performed on a colony-polymerase chain reaction (PCR) amplicon of the V1-V3 region of the 16S rRNA gene (primers 27F and 519R). The MSK strains of *C. nasorum* and *C. yonathiae* were isolated from nasopharyngeal samples collected from infants enrolled in a prospective cohort study that was conducted in Gaborone, Botswana between February 2016 and January 2021 [14]. These samples were inoculated onto plates containing Columbia CNA agar with 5% sheep blood, BHI agar supplemented with 50 µg/mL fosfomycin, and BHI agar with 1% Tween 80 and 50 µg/mL fosfomycin, and incubated aerobically at 37°C in a 5% CO_2_-enriched environment for 48 hours. Preliminary identification of suspected *Corynebacterium* was performed using MALDI-TOF MS or Sanger sequencing on a colony-PCR amplicon of the V1-V3 region of the 16S rRNA gene (primers 27F and 534R). The type strains of *C. accolens* (ATCC 49725^T^) [15], *C. macginleyi* (ATCC 51787^T^) [16], and *C. tuberculostearicum* (ATCC 35692^T^) [10] were obtained from the American Type Culture Collection (Manassas, Virginia).

### Genome Features

Draft genomes of *C. hallux* sp. nov. CTNIH22^T^ and *C. nasorum* sp. nov. KPL3804^T^ were generated through assembly of Illumina sequencing reads, as described previously [12, 17]. Based on four strain genomes, *C. hallux* sp. nov. has an average G+C content of 58.5 mol% and an estimated average genome length of 2.49 Mb, with between 2,293 and 2,427 coding sequences. Based on 13 strain genomes, *C. nasorum* sp. nov. has an average G+C content of 58.5 mol% and an estimated average genome length of 2.46 Mb, with between 2,303 and 2,451 coding sequences (**Table S1**).

### 16S rRNA Gene Phylogeny

PCR amplification of the V1-V9 regions of the 16S rRNA gene of *C. hallux* sp. nov. CTNIH22^T^ and *C. nasorum* sp. nov. KPL3804^T^ was conducted by ACGT, Inc. (Wheeling, IL) and Azenta Life Sciences (South Plainfield, NJ), respectively. Enzymatic cleanup of the PCR products was performed before bidirectional, dye-terminator sequencing on a 3730xl DNA Analyzer (Applied Biosystems, Waltham, MA). For both strains, the corresponding portions of the genome assembly-extracted 16S rRNA gene sequence were determined to be 100% identical to the near-complete gene sequence obtained from PCR amplification and Sanger sequencing. Thus, the full-length genome assembly-extracted 16S rRNA gene sequences were used for subsequent phylogenetic analyses (**Figures 1** and **S1A**).

**Figure 1.**
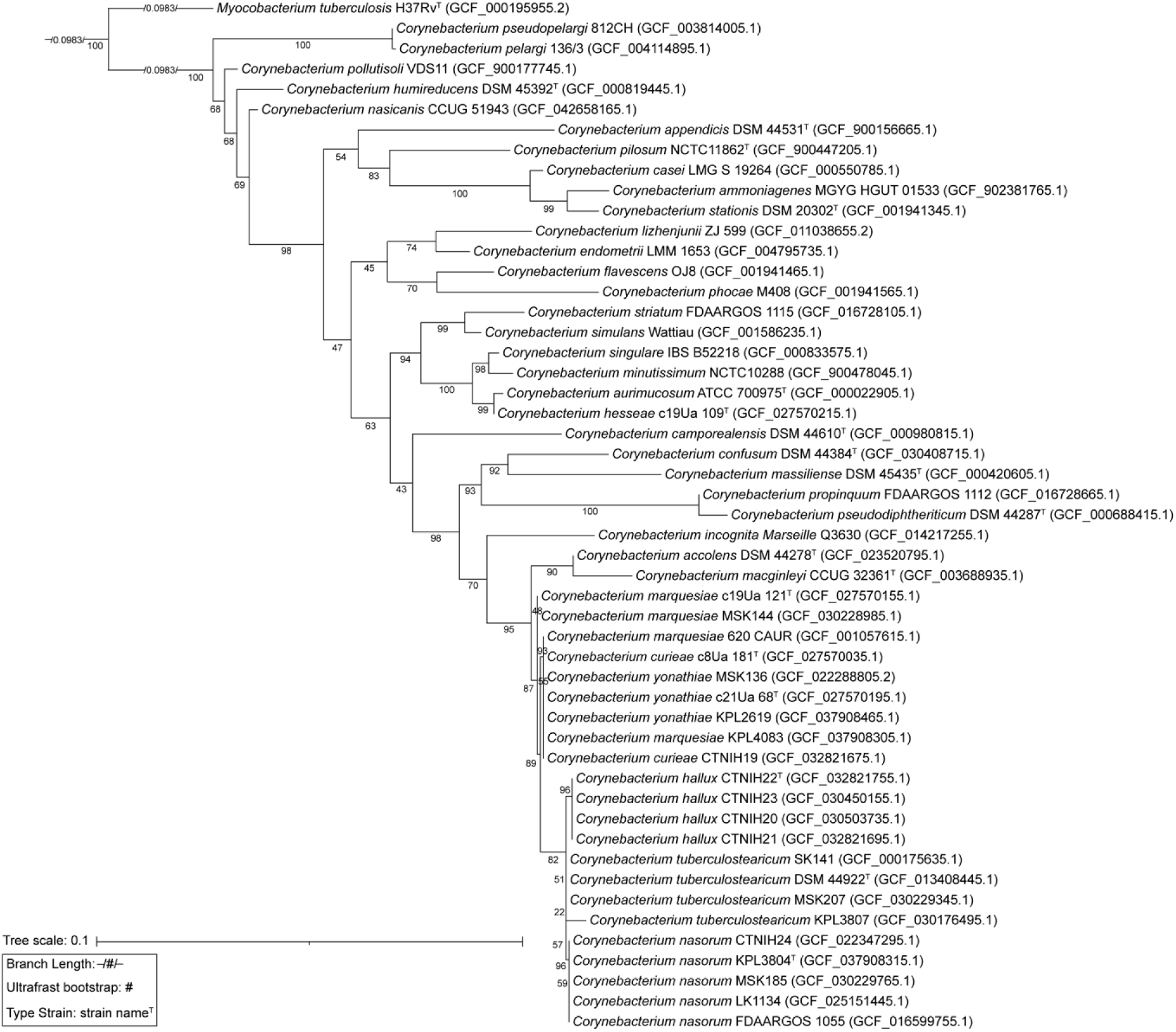
Maximum-likelihood 16S rRNA gene phylogeny of new isolates and type strains of *Corynebacterium* species. A maximum likelihood phylogeny based on nearly full-length 16S rRNA genes of *Corynebacterium* species with ultrafast bootstrap values below 95 at many nodes. *Mycobacterium tuberculosis* H37Rv^T^ is the designated outgroup.

We note that we initially assigned strains of *C. hallux* sp. nov. to the species *C. tuberculostearicum* based on their 16S rRNA gene sequences, and denoted these as ribotype B strains based on differences with the 16S rRNA gene sequences of other known *C. tuberculostearicum* strains [12]. We subsequently used the Type (Strain) Genome Server to determine that genome-sequenced strains of *C. hallux* sp. nov. were most closely related to *C. tuberculostearicum*, although with average nucleotide identity calculations based on the BLAST+ algorithm (ANIb) values of <95% in comparisons to *C. tuberculostearicum* reference genomes [18]. Similarly, we used the Genome Taxonomy Database Toolkit (GTDB-Tk) to determine that *C. nasorum* sp. nov. was closely related to *C. tuberculostearicum* [19]. Also, because the 16S rRNA gene sequences of *C. tuberculostearicum* strain ATCC 35692^T^ and the proposed *C. nasorum* sp. nov. strains are 99.9% identical, it is highly probable that *C. nasorum* sp. nov. sequences were misassigned to *C. tuberculostearicum* in past 16S rRNA gene-based microbiome studies.

A 16S rRNA gene maximum-likelihood phylogeny was constructed using the following species: 1) all of the validly named hits with ≥ 95.7% identity to the 16S rRNA gene of *C. hallux* sp. nov. CTNIH22^T^ and/or *C. nasorum* sp. nov. KPL3804^T^ using the 16S-based ID service on EZBioCloud [20]; 2) additional species that were closely related based on genomes in GTDB-Tk; and 3) *Mycobacterium tuberculosis*^T^ as an outgroup (**Figure 1**). In addition, a larger unrooted 16S rRNA gene phylogeny was constructed to set the two proposed novel species in a broader context within the genus *Corynebacterium* (**Figure S1A**). The 16S rRNA gene-based phylogenies had a large number of poorly supported branches based on ultrafast bootstrap values (**Figures 1** and **S1A**) [21]. This was expected given that the inadequacy of using the 16S rRNA gene alone for constructing reliable phylogenies within the genus *Corynebacterium* is well described [4]. Among *Corynebacterium* species, the *rpoB* gene has more polymorphisms [4], and a phylogeny based on the *rpoB* gene (**Figure S1B**) had a branching pattern with higher support than the 16S rRNA gene-based phylogeny (**Figure S1A**).

### Average Nucleotide Identity

ANIb calculations performed using the Python package pyani v0.2.9 [22, 23] indicated that the genome sequences for the new strains and genomes described in this study were < 95% identical to the type strain of *C. tuberculostearicum,* and to other closely related species (**Figure 2**) [10, 11]. In general, an ANIb threshold of 95-96% accurately represents the boundary between prokaryotic species [24]. Strains of the proposed *C. nasorum* sp. nov. had ANIb values above 95% in comparisons to the genome currently called ‘*Corynebacterium kefirresidentii’* (**Figure 2**, purple box) [25]. However, we were unable to find a type strain bearing this name listed in the publicly available catalogs of major strain repositories. In metagenomic analyses, Kalan and colleagues demonstrate genomic material mapping to the genome called ‘*C. kefirresidentii’* is found on human skin but has higher prevalence and relative abundance in human nasal samples [26]. Based on these findings, and the isolation of a number of nasal strains with ANIb values of >95% to this genome by our laboratories, we agree with the assertion by Kalan and colleagues that the human nasal passages are one of the primary habitats of this species. To reflect this, we propose the species name *Corynebacterium nasorum* sp. nov.. We further propose that strains and genomes previously classified as ‘*C. kefirresidentii’* belong to this proposed novel species (**Table S2**).

**Figure 2.**
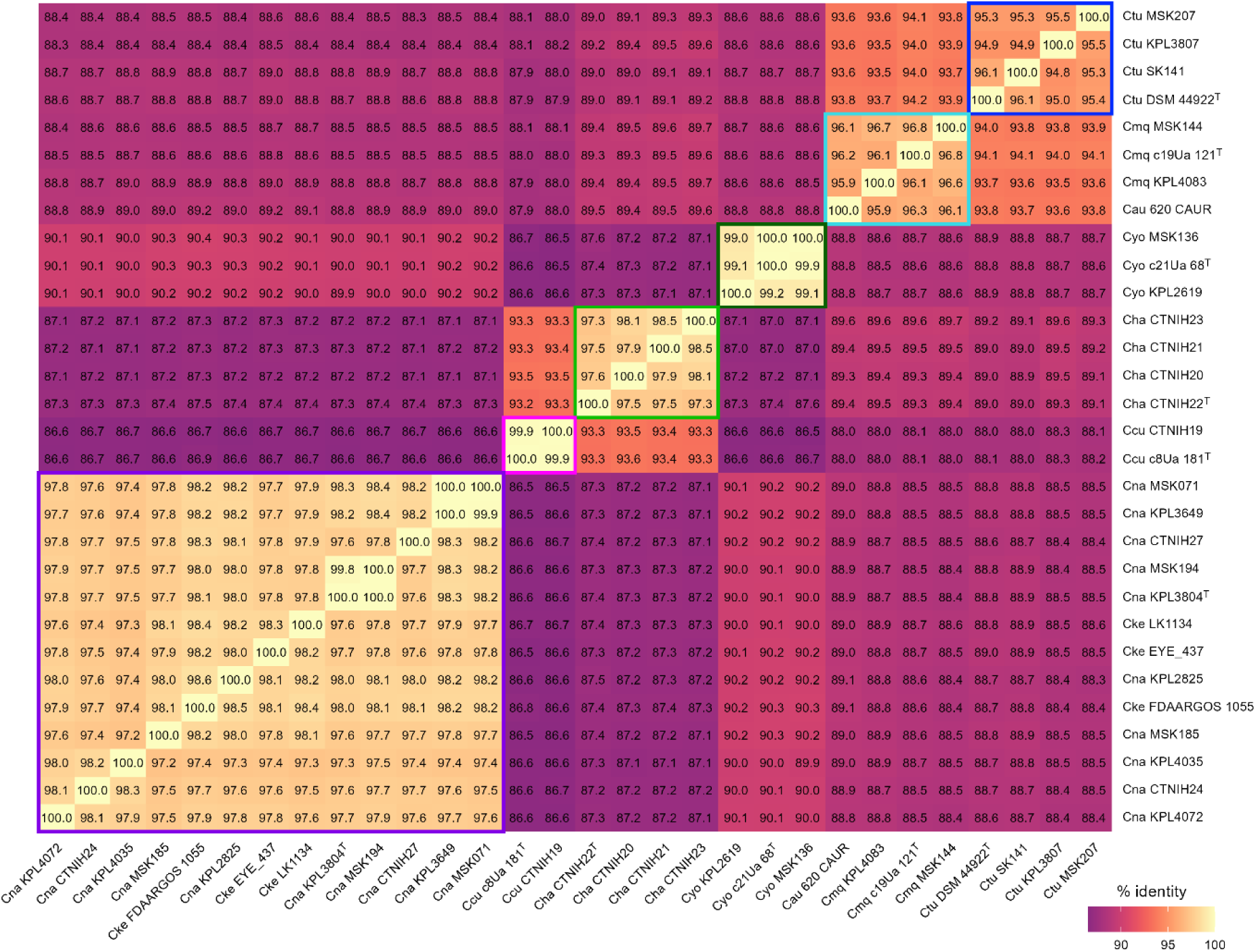
Average nucleotide identity (ANI) among species closely related to *C. tuberculostearicum* revealed two novel species. We used pyani v0.2.9 with ANIb BLAST+ to construct a whole-genome identity ANI heat matrix (see Supplemental Methods) [22, 23]. Boxes highlight species boundaries defined by an ANI threshold of 95% with purple for *C. nasorum* sp. nov. (Cna), pink for *C. curieae* (Ccu), light green for *C. hallux* sp. nov. (Cha), dark green for *C. yonathiae* (Cyo), turquoise for *C. marquesiae* (Cmq), and blue for *C. tuberculostearicum* (Ctu). For *C. tuberculostearicum,* all strains reached a 95% ANI threshold compared to the type strain in at least one direction. The ANIb comparisons indicate that the genome named C. aurimucosum_620_CAUR should be assigned to *C. marquesiae*.

### Phylogenomic Analysis

A maximum-likelihood phylogenomic tree including all of the genome-sequenced strains from Figure 1 with *Mycobacterium tuberculosis* as an outgroup (**Figure 3**) and a phylogenomic tree including the 68 *Corynebacterium* species from Figure S1A (**Figure S1C**) each showed that both *C. hallux* sp. nov. and *C. nasorum* sp. nov. belong to a larger clade that includes *C. tuberculostearicum* and the recently named species *C. curieae*, *C. marquesiae,* and *C. yonathiae* [10, 11]. Kalan and colleagues refer to this monophyletic clade as the “*C. tuberculostearicum* species complex,” and it is most closely related to the clade containing *C. accolens* and *C. macginleyi* (**Figures 3** and **S1C**) [26]. Of note, the genomes currently labeled in GTDB-Tk [19] as “*C. aurimucosum*_E” are misassigned at the species level based on the phylogenomic analyses shown here and those performed by Kalan and colleagues [26], since *C. aurimucosum_*E 620_CAUR [27] clusters far from the genome for the type strain *C. aurimucosum* DSM 44827 (**Figures 3** and **S1C**). Based on an ANIb threshold of 95%, genomes labeled in GTDB-Tk as “*C. aurimucosum*_E” assign to the recently named species *C. marquesiae* (**Figures 2** and **S2A**) [11].

**Figure 3.**
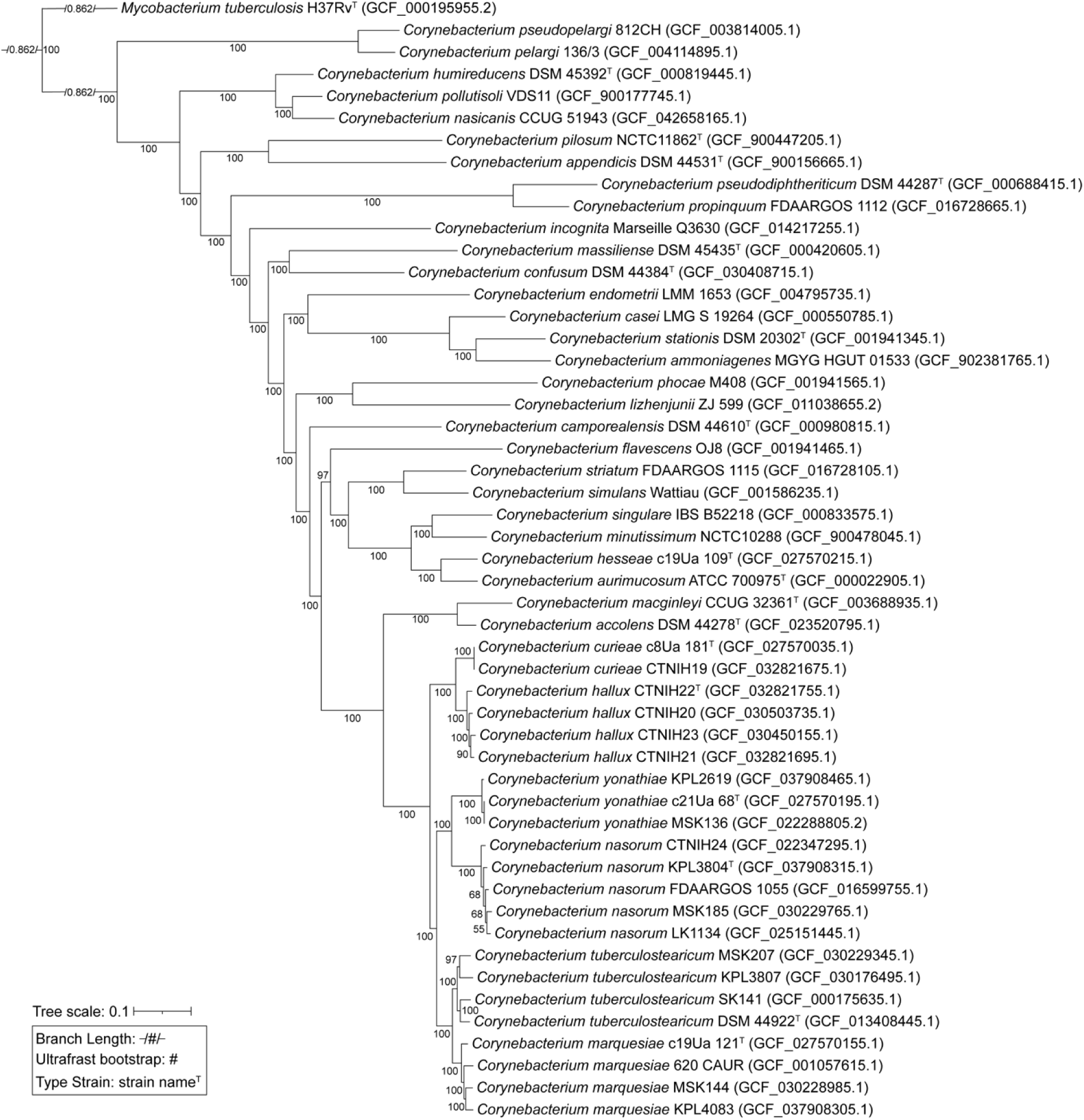
Maximum-likelihood phylogenomic tree of new isolates and type strains of *Corynebacterium* species. A maximum likelihood phylogenomic tree was constructed using 305 shared conservative core gene clusters and *Mycobacterium tuberculosis* H37Rv^T^ as the designated outgroup. This monophyletic tree shows robust separation of *Corynebacterium* species based on ultrafast bootstrap values.

Together, ANIb calculations and the phylogenomic trees confirm that the strains labeled *C. hallux* sp. nov. and *C. nasorum* sp. nov. represent novel species belonging to the genus *Corynebacterium*.

### Comparative Genomic Analysis

The metabolic capabilities of more divergent *Corynebacterium* species sharing the common habitat of the human nasal passages are highly conserved [28]. Although members of the *C. tuberculostearicum* species complex are known to inhabit different human body site habitats including skin [12], the nasal passages [26], and the female urinary tract [11], we hypothesized that these would exhibit conserved metabolic capabilities based on their close phylogenetic relationship to each other (**Figures 3, S1C,** and **S2A**). Indeed, metabolic estimation on genomes of these species using the anvi-run-kegg-kofams and the anvi-estimate-metabolism programs of anvi’o v8 [29, 30], which rely on Kyoto Encyclopedia of Genes and Genomes (KEGG) metabolic annotations [31], revealed largely shared metabolic capabilities with some strain-level variation within specific species (**Figure 4**, **Table S3**; see https://klemonlab.github.io/NovCor_Manuscript/Methods_Anvio.html for detailed methods). This analysis estimated these 30 strain genomes covering six species (**Figures 2** and **S2**) all shared 48 stepwise complete KEGG modules, with most strains also sharing an additional six complete KEGG modules (**Table 1**). These included many modules for amino acid biosynthesis, which is typical of *Corynebacterium* species. We estimated that all 30 strain genomes also encode a complete tricarboxylic acid cycle, consistent with their preference for aerobic growth, along with a number of other modules involved in central carbohydrate metabolism (**Table 1**).

**Figure 4.**
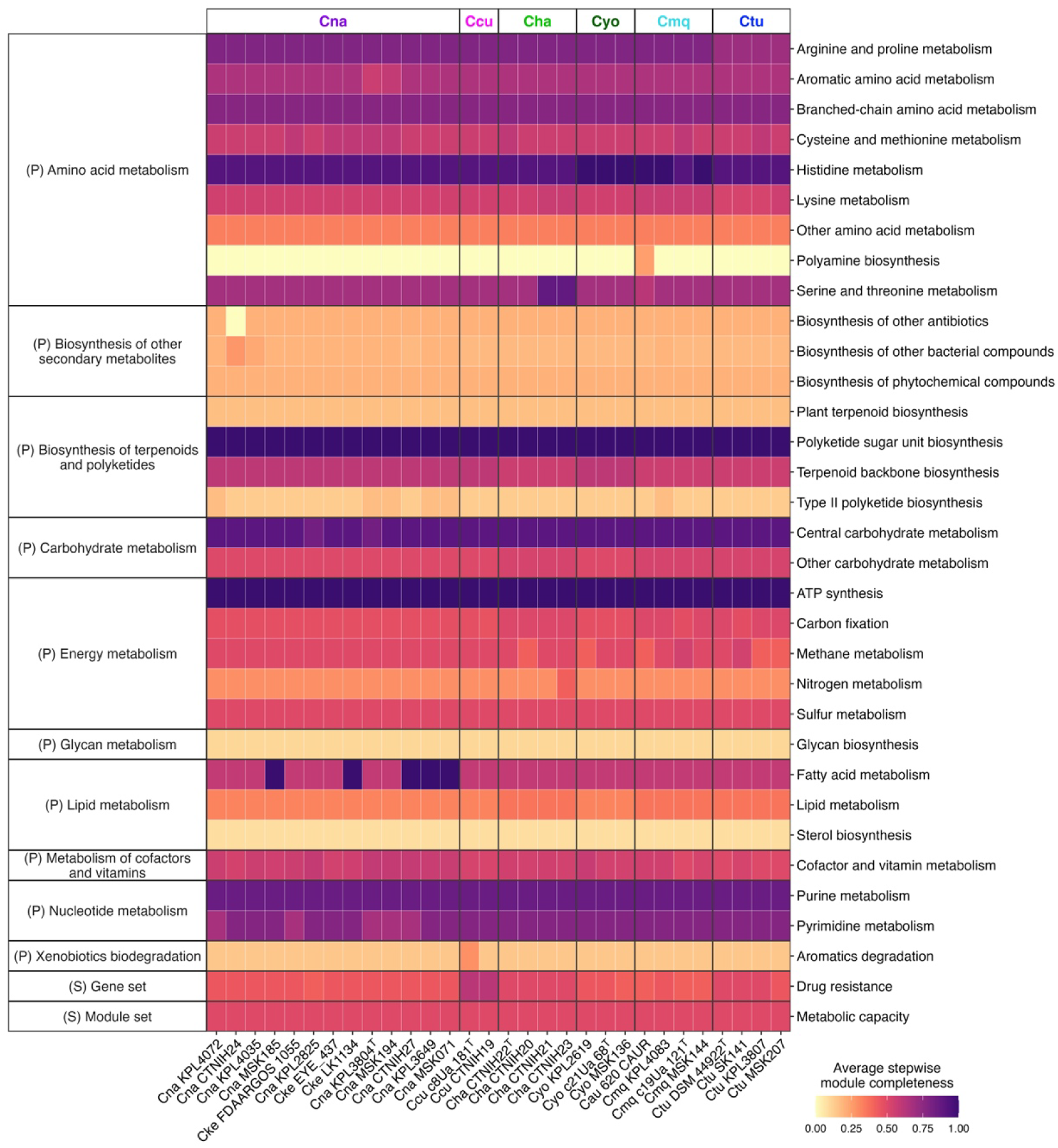
Species closely related to *C. tuberculostearicum* are estimated to largely share a common set of metabolic capabilities. The heatmap represents average estimated module stepwise completion scores by KEGG subcategories for each of the 30 genomes from Figure 2 covering six species that clade closely with *C. tuberculostearicum* (Figures 3 **and S2A**). Average stepwise completion scores were calculated including only modules detected in at least one of the analyzed genomes. (P) represents pathway modules; (S) represents signature modules. Cna, *C. nasorum* sp. nov.; Ccu, *C. curieae*; Cha, *C. hallux* sp. nov.; Cyo, *C. yonathiae*; Cmq, *C. marquesiae*; Ctu, *C. tuberculostearicum*.

**Table 1.** Estimated stepwise complete KEGG modules shared by at least 26 of the 30. *Corynebacterium* strain genomes belonging to species closely related to *C. tuberculostearicum,* including *C. nasorum* sp. nov. and *C. hallux* sp. nov. (# represents number of genomes)

| Module ID | Module Name | Module Subcategory | # |
| --- | --- | --- | --- |
| M00015 | Proline biosynthesis, glutamate => proline | Arginine and proline metabolism | 30 |
| M00844 | Arginine biosynthesis, ornithine => arginine | Arginine and proline metabolism | 30 |
| M00970 | Proline degradation, proline => glutamate | Arginine and proline metabolism | 30 |
| M00022 | Shikimate pathway, phosphoenolpyruvate + erythrose-4P => chorismite | Aromatic amino acid metabolism | 30 |
| M00019 | Valine/isoleucine biosynthesis, pyruvate => valine / 2-oxobutanoate => isoleucine | Branched-chain amino acid metabolism | 30 |
| M00432 | Leucine biosynthesis, 2-oxoisovalerate => 2-oxoisocaproate | Branched-chain amino acid metabolism | 30 |
| M00570 | Isoleucine biosynthesis, threonine => 2-oxobutanoate => isoleucine | Branched-chain amino acid metabolism | 30 |
| M00017 | Methionine biosynthesis, aspartate => homoserine => methionine | Cysteine and methionine metabolism | 30 |
| M00021 | Cysteine biosynthesis, serine => cysteine | Cysteine and methionine metabolism | 30 |
| M00045 | Histidine degradation, histidine => N-formiminoglutamate => glutamate | Histidine metabolism | 30 |
| M00018 | Threonine biosynthesis, aspartate => homoserine => threonine | Serine and threonine metabolism | 30 |
| M00621 | Glycine cleavage system | Serine and threonine metabolism | 30 |
| M00793 | dTDP-L-rhamnose biosynthesis | Polyketide sugar unit biosynthesis | 30 |
| M00096 | C5 isoprenoid biosynthesis, non-mevalonate pathway | Terpenoid backbone biosynthesis | 30 |
| M00364 | C10-C20 isoprenoid biosynthesis, bacteria | Terpenoid backbone biosynthesis | 30 |
| M00365 | C10-C20 isoprenoid biosynthesis, archaea | Terpenoid backbone biosynthesis | 30 |
| M00001 | Glycolysis (Embden-Meyerhof pathway), glucose => pyruvate | Central carbohydrate metabolism | 30 |
| M00002 | Glycolysis, core module involving three-carbon compounds | Central carbohydrate metabolism | 30 |
| M00003 | Gluconeogenesis, oxaloacetate => fructose-6P | Central carbohydrate metabolism | 30 |
| M00004 | Pentose phosphate pathway (Pentose phosphate cycle) | Central carbohydrate metabolism | 30 |
| M00005 | PRPP biosynthesis, ribose 5P => PRPP | Central carbohydrate metabolism | 30 |
| M00006 | Pentose phosphate pathway, oxidative phase, glucose 6P => ribulose 5P | Central carbohydrate metabolism | 30 |
| M00007 | Pentose phosphate pathway, non-oxidative phase, fructose 6P => ribose 5P | Central carbohydrate metabolism | 30 |
| M00009 | Citrate cycle (TCA cycle, Krebs cycle) | Central carbohydrate metabolism | 30 |
| M00010 | Citrate cycle, first carbon oxidation, oxaloacetate => 2-oxoglutarate | Central carbohydrate metabolism | 30 |
| M00011 | Citrate cycle, second carbon oxidation, 2-oxoglutarate => oxaloacetate | Central carbohydrate metabolism | 30 |
| M00549 | Nucleotide sugar biosynthesis, glucose => UDP-glucose | Other carbohydrate metabolism | 30 |
| M00554 | Nucleotide sugar biosynthesis, galactose => UDP-galactose | Other carbohydrate metabolism | 30 |
| M00632 | Galactose degradation, Leloir pathway, galactose => alpha-D-glucose-1P | Other carbohydrate metabolism | 30 |
| M00909 | UDP-N-acetyl-D-glucosamine biosynthesis, prokaryotes, glucose => UDP-GlcNAc | Other carbohydrate metabolism | 30 |
| M00151 | Cytochrome bc1 complex respiratory unit | ATP synthesis | 30 |
| M00155 | Cytochrome c oxidase, prokaryotes | ATP synthesis | 30 |
| M00157 | F-type ATPase, prokaryotes and chloroplasts | ATP synthesis | 30 |
| M00579 | Phosphate acetyltransferase-acetate kinase pathway, acetyl-CoA => acetate | Carbon fixation | 30 |
| M00086 | beta-Oxidation, acyl-CoA synthesis | Fatty acid metabolism | 30 |
| M00120 | Coenzyme A biosynthesis, pantothenate => CoA | Cofactor and vitamin metabolism | 30 |
| M00121 | Heme biosynthesis, plants and bacteria, glutamate => heme | Cofactor and vitamin metabolism | 30 |
| M00125 | Riboflavin biosynthesis, plants and bacteria, GTP => riboflavin/FMN/FAD | Cofactor and vitamin metabolism | 30 |
| M00126 | Tetrahydrofolate biosynthesis, GTP => THF | Cofactor and vitamin metabolism | 30 |
| M00881 | Lipoic acid biosynthesis, plants and bacteria, octanoyl-ACP => dihydrolipoyl-E2/H | Cofactor and vitamin metabolism | 30 |
| M00899 | Thiamine salvage pathway, HMP/HET => TMP | Cofactor and vitamin metabolism | 30 |
| M00916 | Pyridoxal-P biosynthesis, R5P + glyceraldehyde-3P + glutamine => pyridoxal-P | Cofactor and vitamin metabolism | 30 |
| M00926 | Heme biosynthesis, bacteria, glutamyl-tRNA => coproporphyrin III => heme | Cofactor and vitamin metabolism | 30 |
| M00048 | De novo purine biosynthesis, PRPP + glutamine => IMP | Purine metabolism | 30 |
| M00049 | Adenine ribonucleotide biosynthesis, IMP => ADP,ATP | Purine metabolism | 30 |
| M00050 | Guanine ribonucleotide biosynthesis, IMP => GDP,GTP | Purine metabolism | 30 |
| M00053 | Deoxyribonucleotide biosynthesis, ADP/GDP/CDP/UDP => dATP/dGTP/dCTP/dUTP | Purine metabolism | 30 |
| M00938 | Pyrimidine deoxyribonucleotide biosynthesis, UDP => dTTP | Pyrimidine metabolism | 30 |
| M00023 | Tryptophan biosynthesis, chorismate => tryptophan | Aromatic amino acid metabolism | 29 |
| M00020 | Serine biosynthesis, glycerate-3P => serine | Serine and threonine metabolism | 29 |
| M00140 | C1-unit interconversion, prokaryotes | Cofactor and vitamin metabolism | 29 |
| M00307 | Pyruvate oxidation, pyruvate => acetyl-CoA | Central carbohydrate metabolism | 28 |
| M00028 | Ornithine biosynthesis, glutamate => ornithine | Arginine and proline metabolism | 26 |
| M00016 | Lysine biosynthesis, succinyl-DAP pathway, aspartate => lysine | Lysine metabolism | 26 |

### Phenotypic and Chemotaxonomic Characterisation

Growth of strains included in this study was first determined on the following solid media: tryptic soy agar (TSA) with 5% sheep blood (Remel, Lenexa, KS), BHI agar (Becton Dickinson, Franklin Lakes, NJ), and BHI with 1% Tween 80 (MilliporeSigma, Burlington, MA). Isolates were suspended in sterile phosphate buffered saline (Genesee Scientific, El Cajon, CA) to an OD_600_ of 0.10–0.15 and plated in a quadrant pattern using a sterile 10-µl loop. Growth was judged by two authors (EBP, MSK) based on the size and density of colonies on these plates. Given that all strains grew well on BHI with 1% Tween 80 plates, growth on this medium was further evaluated for up to 14 days at various temperature (4°C, 20°C, 30°C, 37°C, 42°C, 50°C) and atmospheric (aerobic/5% CO_2_, microaerophilic, anaerobic) conditions. Microaerophilic and anaerobic conditions were generated using the AnaeroPack system with MicroAero and Anaerobic gas generators (Thermo Fisher Scientific, Waltham, MA). Growth of all strains was observed at temperatures between 30°C and 42°C, with *C. accolens* ATCC 49725^T^, *C. tuberculostearicum* ATCC 35692^T^, *C. hallux* sp. nov. CTNIH22^T^, and strains of *C. yonathiae* additionally demonstrating growth at 20°C (**Table 2**). Growth was also observed for all strains in microaerophilic conditions, with *C. accolens* ATCC 49725^T^, *C. macginleyi* ATCC 51787^T^, and strains of *C. nasorum* sp. nov. and *C. yonathiae* additionally demonstrating weak growth in anaerobic conditions (**Table 2**).

**Table 2.**
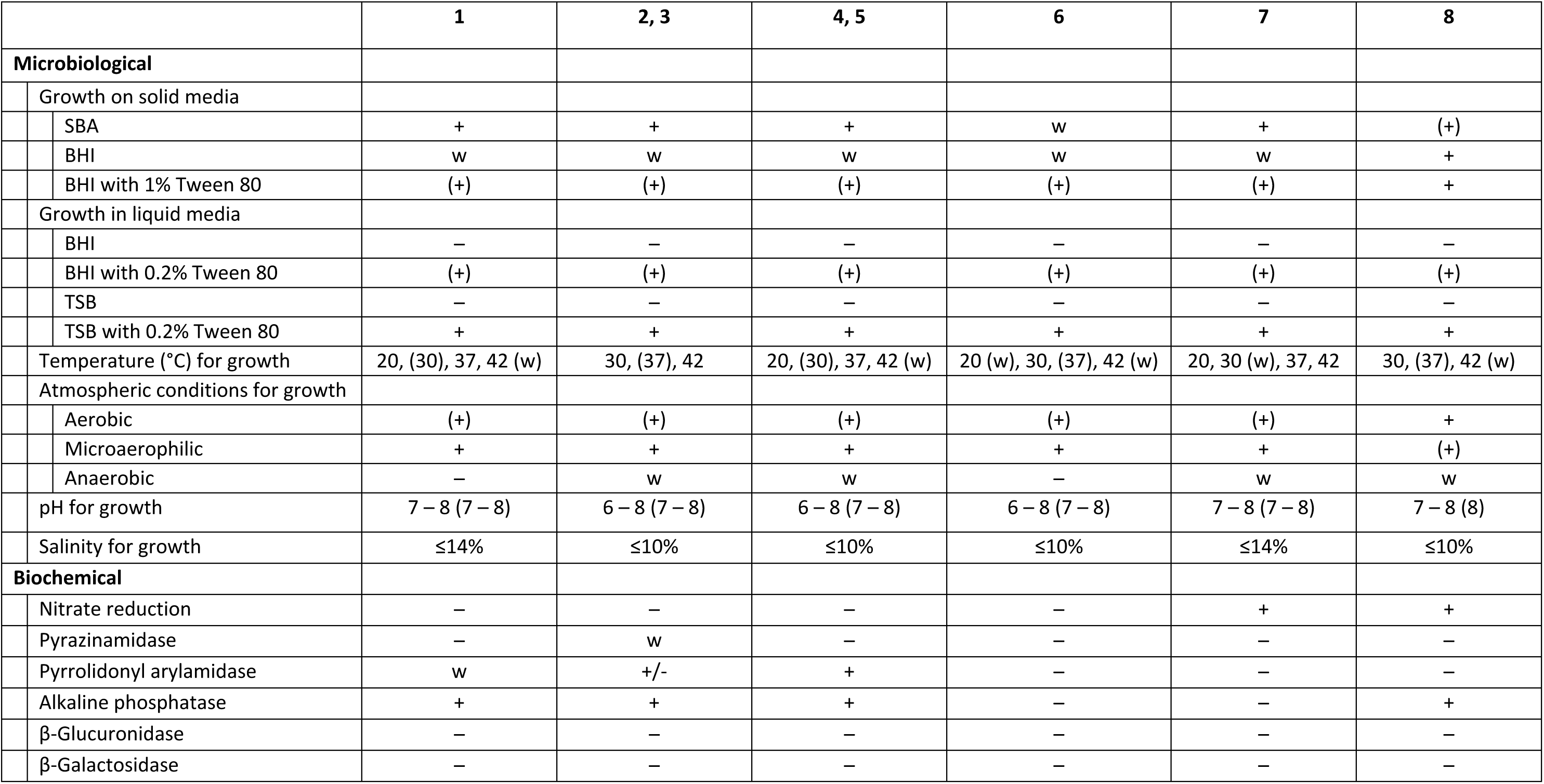
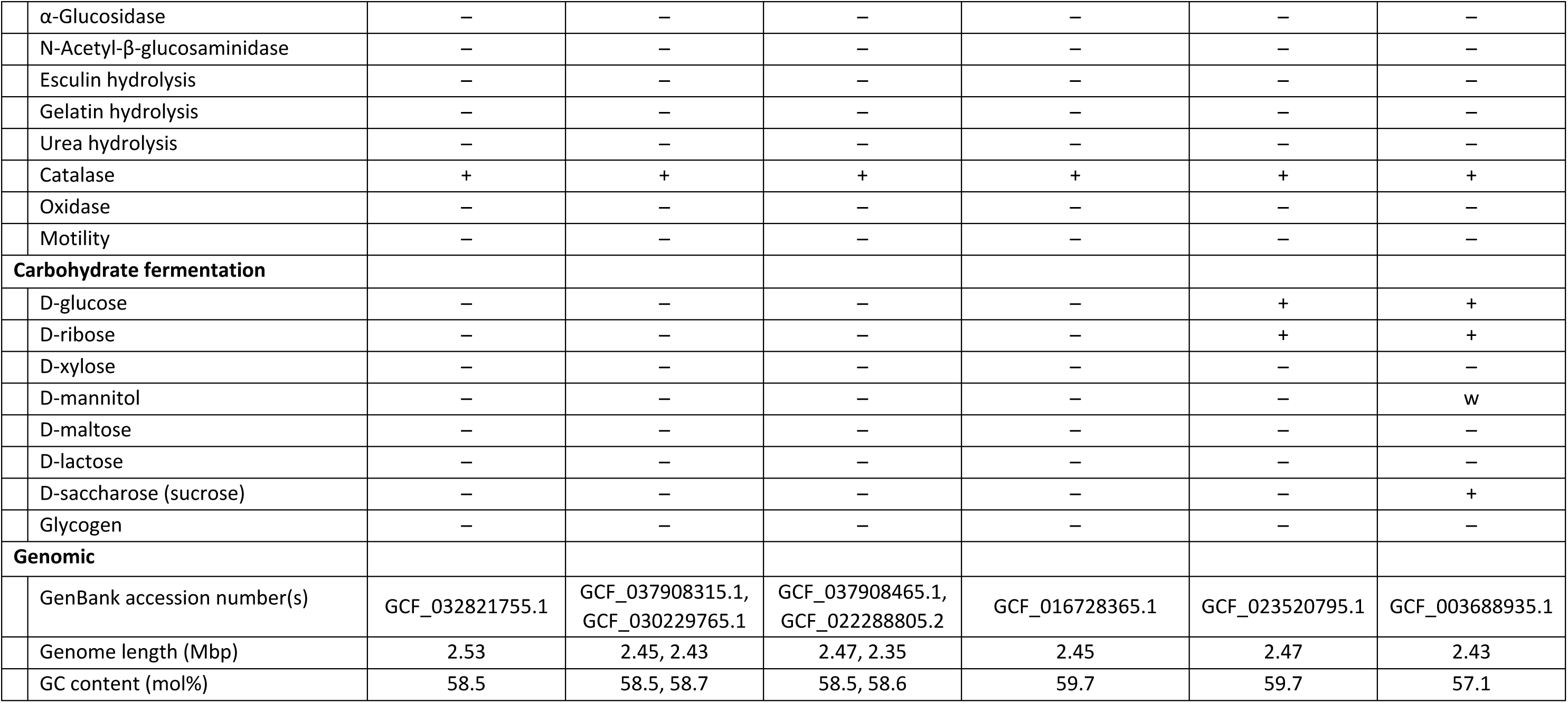
Microbiological, biochemical, and genomic characteristics of *Corynebacterium* strains. Strains: 1, *C. hallux* sp. nov. CTNIH22^T^; 2, *C. nasorum* sp. nov. KPL3804^T^; 3, *C. nasorum* sp. nov. MSK185; 4, *C. yonathiae* MSK136; 5, *C. yonathiae* KPL2619; 6, *C. tuberculostearicum* ATCC 35692^T^; 7, *C. accolens* ATCC 49725^T^; 8, *C. macginleyi* ATCC 51787^T^. Data were generated in this study for all strains. +, positive; w, weakly positive; -, negative; +/-, variable. BHI, brain-heart infusion; SBA, tryptic soy agar with 5% sheep blood; TSB, tryptic soy broth. Results in parentheses indicate optimal values.

|  | 1 | 2, 3 | 4, 5 | 6 | 7 | 8 |
| --- | --- | --- | --- | --- | --- | --- |
| **Microbiological** |  |  |  |  |  |  |
| Growth on solid media |  |  |  |  |  |  |
| SBA | + | + | + | w | + | (+) |
| BHI | w | w | w | w | w | + |
| BHI with 1% Tween 80 | (+) | (+) | (+) | (+) | (+) | + |
| Growth in liquid media |  |  |  |  |  |  |
| BHI | – | – | – | – | – | – |
| BHI with 0.2% Tween 80 | (+) | (+) | (+) | (+) | (+) | (+) |
| TSB | – | – | – | – | – | – |
| TSB with 0.2% Tween 80 | + | + | + | + | + | + |
| Temperature (°C) for growth | 20, (30), 37, 42 (w) | 30, (37), 42 | 20, (30), 37, 42 (w) | 20 (w), 30, (37), 42 (w) | 20, 30 (w), 37, 42 | 30, (37), 42 (w) |
| Atmospheric conditions for growth |  |  |  |  |  |  |
| Aerobic | (+) | (+) | (+) | (+) | (+) | + |
| Microaerophilic | + | + | + | + | + | (+) |
| Anaerobic | – | w | w | – | w | w |
| pH for growth | 7 – 8 (7 – 8) | 6 – 8 (7 – 8) | 6 – 8 (7 – 8) | 6 – 8 (7 – 8) | 7 – 8 (7 – 8) | 7 – 8 (8) |
| Salinity for growth | ≤14% | ≤10% | ≤10% | ≤10% | ≤14% | ≤10% |
| **Biochemical** |  |  |  |  |  |  |
| Nitrate reduction | – | – | – | – | + | + |
| Pyrazinamidase | – | w | – | – | – | – |
| Pyrrolidonyl arylamidase | w | +/- | + | – | – | – |
| Alkaline phosphatase | + | + | + | – | – | + |
| β-Glucuronidase | – | – | – | – | – | – |
| β-Galactosidase | – | – | – | – | – | – |

|  |  |  |  |  |  |  |  |
| --- | --- | --- | --- | --- | --- | --- | --- |
| | α-Glucosidase | – | – | – | – | – | – |
| | N-Acetyl-β-glucosaminidase | – | – | – | – | – | – |
|  | Esculin hydrolysis | – | – | – | – | – | – |
|  | Gelatin hydrolysis | – | – | – | – | – | – |
|  | Urea hydrolysis | – | – | – | – | – | – |
|  | Catalase | + | + | + | + | + | + |
|  | Oxidase | – | – | – | – | – | – |
|  | Motility | – | – | – | – | – | – |
| **Carbohydrate fermentation** |  |  |  |  |  |  |  |
|  | D-glucose | – | – | – | – | + | + |
|  | D-ribose | – | – | – | – | + | + |
|  | D-xylose | – | – | – | – | – | – |
|  | D-mannitol | – | – | – | – | – | w |
|  | D-maltose | – | – | – | – | – | – |
|  | D-lactose | – | – | – | – | – | – |
|  | D-saccharose (sucrose) | – | – | – | – | – | + |
|  | Glycogen | – | – | – | – | – | – |
| **Genomic** |  |  |  |  |  |  |  |
|  | GenBank accession number(s) | GCF_032821755.1 | GCF_037908315.1,<br>GCF_030229765.1 | GCF_037908465.1,<br>GCF_022288805.2 | GCF_016728365.1 | GCF_023520795.1 | GCF_003688935.1 |
|  | Genome length (Mbp) | 2.53 | 2.45, 2.43 | 2.47, 2.35 | 2.45 | 2.47 | 2.43 |
|  | GC content (mol%) | 58.5 | 58.5, 58.7 | 58.5, 58.6 | 59.7 | 59.7 | 57.1 |

*C. hallux* sp. nov. CTNIH22^T^ typically grew as creamy, white colonies measuring 3-5 mm in diameter on BHI with 1% Tween 80 agar; growth was weaker on TSA with 5% sheep blood or on BHI agar without Tween 80, with colonies measuring 1-2 mm in diameter that were non-hemolytic on blood-containing agar. *C. nasorum* sp. nov. strains grew optimally on BHI with 1% Tween 80 agar, yielding large white colonies 5-10 mm in diameter. Colonies of *C. nasorum* sp. nov. were non-hemolytic on TSA with 5% sheep blood agar. *C. yonathiae* strains appeared as raised, creamy colonies between 5-10 mm in diameter when grown on BHI with 1% Tween 80 agar. Colonies on TSA with 5% sheep blood were flat, translucent, non-hemolytic, and approximately 2-3 mm in diameter.

Growth in liquid media was assessed using BHI broth, BHI broth with 0.2% Tween 80, tryptic soy broth (TSB), and TSB with 0.2% Tween 80. For these assays, culture tubes were inoculated with bacterial cells washed twice with phosphate buffered saline to remove traces of Tween 80 retained from the solid media, then resuspended to an OD_600_ of 0.10–0.15. All cultures reached an OD_600_ at or above 2.0 at 48 hours after inoculation into BHI broth with 0.2% Tween 80. Therefore, this liquid medium was used for subsequent assays testing for growth at varying pH and (2.0, 4.0, 6.0, 7.0, 8.0, 10.0, 12.0) and salinity (0%, 3%, 5%, 7%, 10%, 14%, 20%). Liquid culture tubes were incubated aerobically at 37°C with shaking at 200 rpm for 72 hours or until the cultures exceeded an OD_600_ of 2.0. The pH range for growth of most strains was 7–8, with optimal growth observed at pH 8 (**Table 2**). Growth of all strains was observed at salinity at or below 10%, with *C. accolens* ATCC 49725^T^ and *C. hallux* sp. nov. CTNIH22^T^ demonstrating growth at salinity up to 14% (**Table 2**). All strains also demonstrated growth in TSB with 0.2% Tween 80. No growth was observed in BHI or TSB broth in the absence of Tween 80.

Gram stains were performed using fresh cultures grown on BHI with 1% Tween 80 agar and using a commercial kit (Hardy Diagnostics, Santa Maria, CA). Cells of *C. hallux* sp. nov. CTNIH^T^ and strains of *C. nasorum* sp. nov. and *C. yonathiae* were Gram-positive, non-spore-forming irregular rods or coccoid. For visualization by scanning electron microscopy, cells were fixed with a solution containing 2% glutaraldehyde and 4% formaldehyde prior to transfer to the Duke University Shared Materials Instrumentation Facility. In scanning electron microscopy images, *C. hallux* sp. nov. cells **Figure 5A**) appeared as pleomorphic rods to coccoid, with most cells being 0.6-2.0 µm long and 0.4-0.6 µm wide. *C. nasorum* sp. nov. cells (**Figure 5B**) similarly appeared as heterogenous rods to coccoid, with most cells measuring between 0.6-2.2 µm long and 0.4-0.6 µm wide. *C. yonathiae* cells (**Figure 5C**) were coccoid to elongated rods measuring up to 5 µm in length and 0.4-0.6 µm wide.

**Figure 5.**
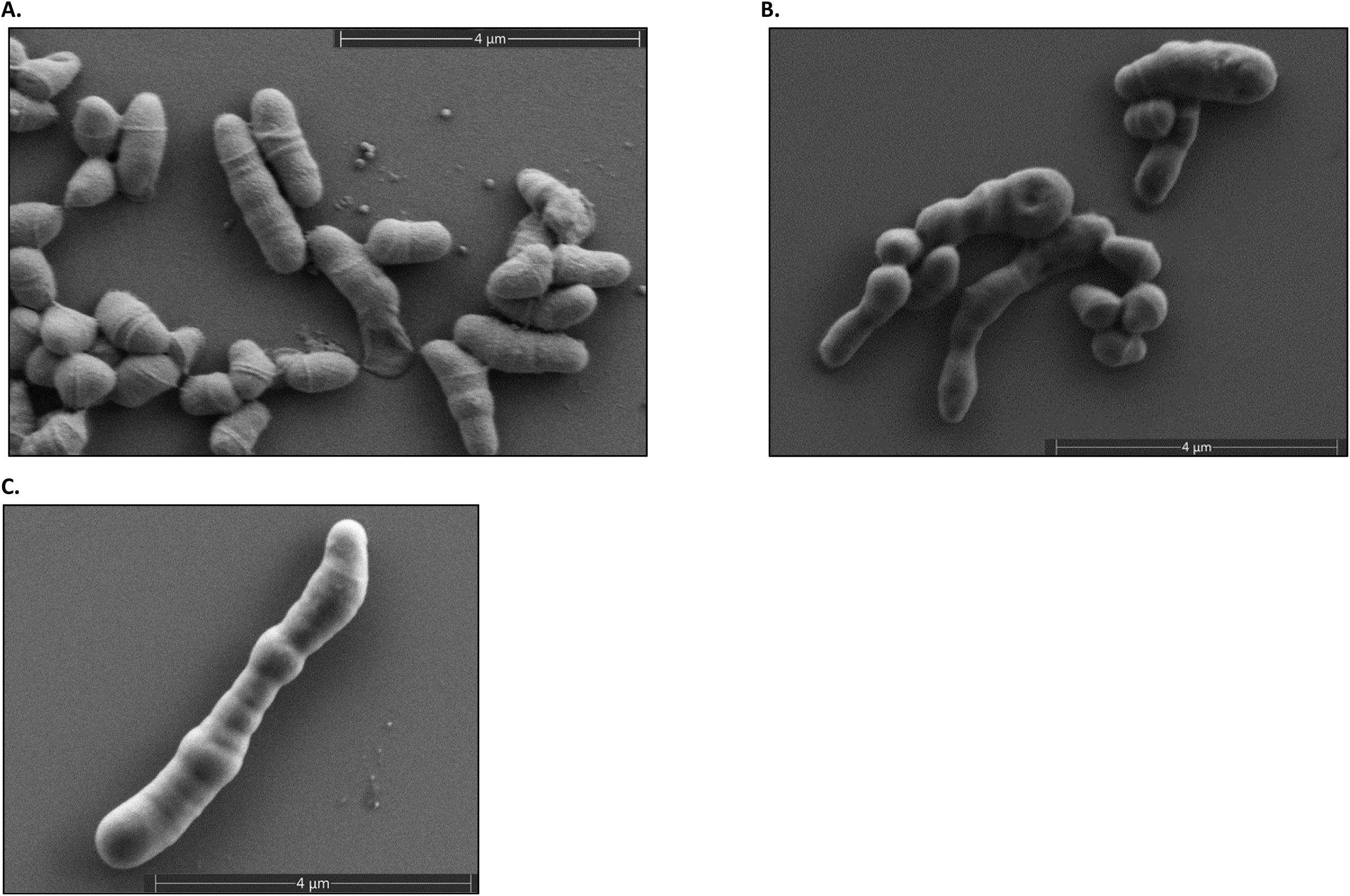
Scanning electron microscopy images (12,000x) of strains *C. hallux* sp. nov. CTNIH22^T^ (**A**), *C. nasorum* sp. nov. KPL3804^T^ (**B**), and *C. yonathiae* KPL2619 (**C**).

Enzymatic (including catalase) and fermentation activities were tested using API CORYNE strips (bioMérieux, Marcy-l‘Etoile, France) according to the manufacturer’s instructions. Oxidase testing was performed using OxiStrips (Hardy Diagnostics, Santa Maria, CA) according to the package insert. Motility was assessed by stabbing culture tubes containing BHI with 1% Tween 80 and 0.5% agar with a fresh culture of each *Corynebacterium* strain; *Pseudomonas aeruginosa, Escherichia coli*, and *Staphylococcus aureus* were used as comparators for this assay. All strains were catalase-positive, oxidase-negative, and non-motile (**Table 3**). On biochemical testing, positive reactions for alkaline phosphatase were observed for *C. macginleyi* ATCC 51787^T^, *C. hallux* sp. nov. CTNIH22^T^, and strains of *C. nasorum* sp. nov. and *C. yonathiae* (**Table 3**). *C. hallux* sp. nov. CTNIH22^T^ and strains of *C. yonathiae* sp. nov. had a positive reaction for pyrrolidonyl arylamidase, while this activity was variable for strains of *C. nasorum* sp. nov. Nitrate reduction was observed for *C. accolens* ATCC 49725^T^ and *C. macginleyi* ATCC 51787^T^, while strains of *C. nasorum* sp. nov. had a weakly positive reaction for pyrazinamidase. Fermentation of D-glucose and D-ribose was noted for strains *C. accolens* ATCC 49725^T^ and *C. macginleyi* ATCC 51787^T^, while *C. macginleyi* ATCC 51787^T^ additionally fermented D-mannitol and D-saccharose (**Table 3**). No carbohydrate fermentation was noted for *C. hallux* sp. nov. CTNIH22^T^ or strains of *C. nasorum* sp. nov. or *C. yonathiae*.

**Table 3.**
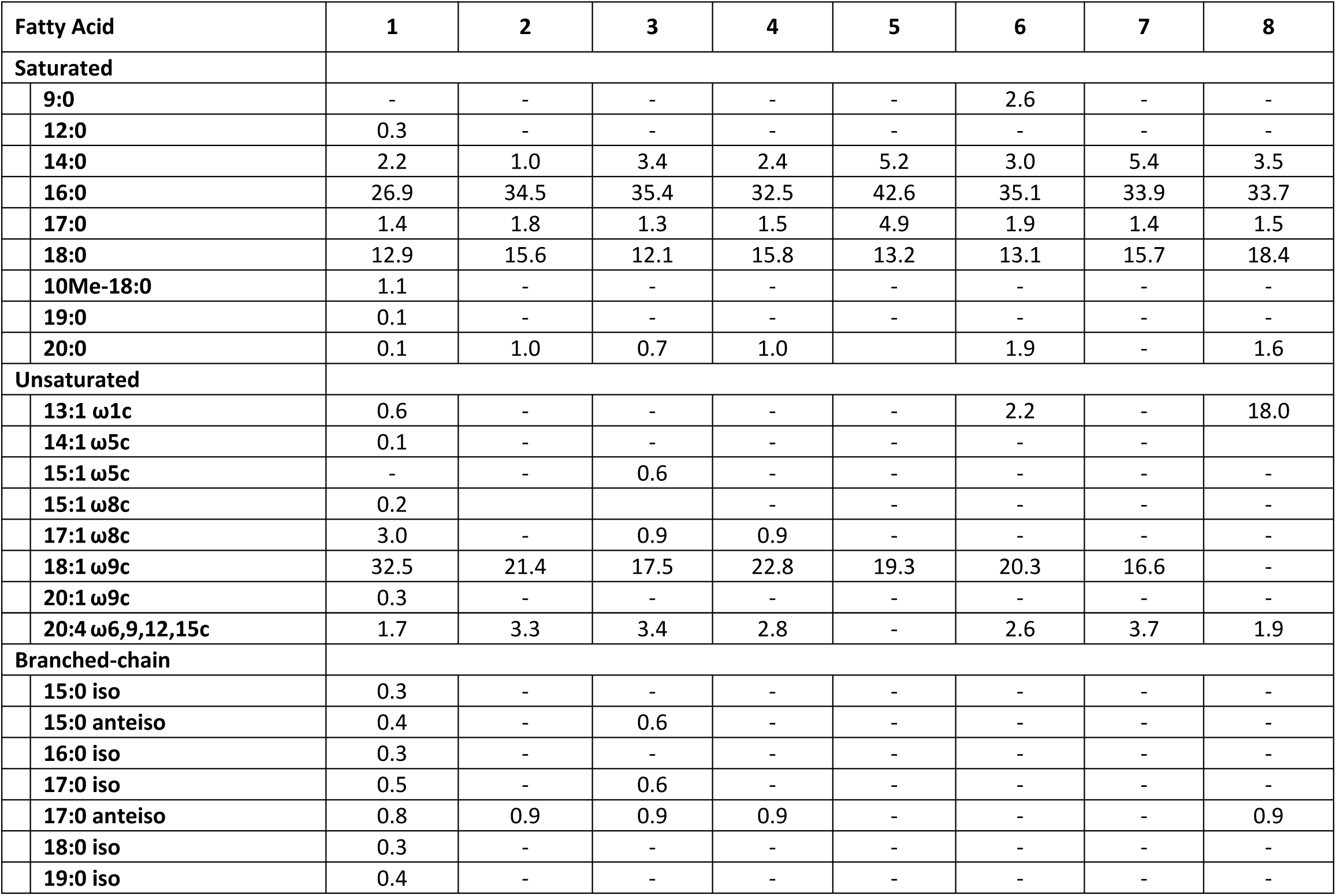
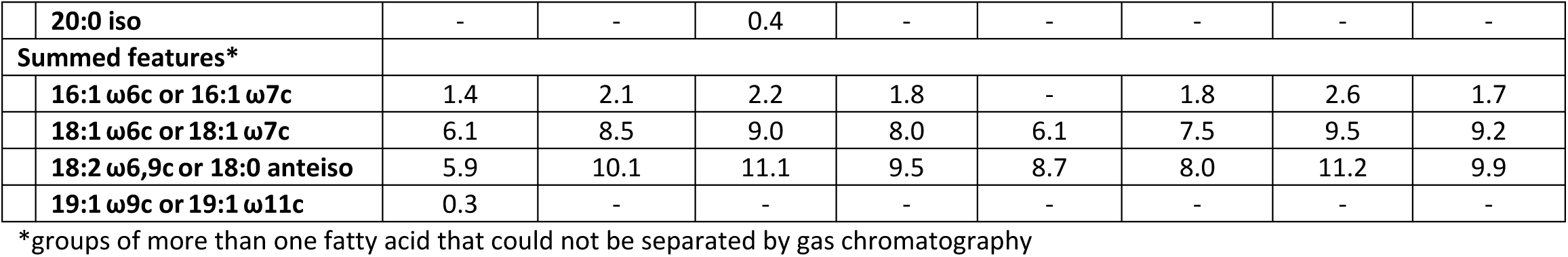
Cellular fatty acid composition of *Corynebacterium* spp. strains by fatty acid methyl esters (FAME) analysis. Strains: 1, *C. hallux* sp. nov. CTNIH22^T^; 2, *C. nasorum* sp. nov. KPL3804^T^; 3, *C. nasorum* sp. nov. MSK185; 4, *C. yonathiae* MSK136; 5, *C. yonathiae* KPL2619; 6, *C. tuberculostearicum* ATCC 35692^T^; 7, *C. accolens* ATCC 49725^T^; 8, *C. macginleyi* ATCC 51787^T^. Data were generated in this study for all strains. –, not detected. Percentages may not sum to 100% due to rounding.

| Fatty Acid | 1 | 2 | 3 | 4 | 5 | 6 | 7 | 8 |
| --- | --- | --- | --- | --- | --- | --- | --- | --- |
| **Saturated** |  |  |  |  |  |  |  |  |
| 9:0 | - | - | - | - | - | 2.6 | - | - |
| 12:0 | 0.3 | - | - | - | - | - | - | - |
| 14:0 | 2.2 | 1.0 | 3.4 | 2.4 | 5.2 | 3.0 | 5.4 | 3.5 |
| 16:0 | 26.9 | 34.5 | 35.4 | 32.5 | 42.6 | 35.1 | 33.9 | 33.7 |
| 17:0 | 1.4 | 1.8 | 1.3 | 1.5 | 4.9 | 1.9 | 1.4 | 1.5 |
| 18:0 | 12.9 | 15.6 | 12.1 | 15.8 | 13.2 | 13.1 | 15.7 | 18.4 |
| 10Me-18:0 | 1.1 | - | - | - | - | - | - | - |
| 19:0 | 0.1 | - | - | - | - | - | - | - |
| 20:0 | 0.1 | 1.0 | 0.7 | 1.0 |  | 1.9 | - | 1.6 |
| **Unsaturated** |  |  |  |  |  |  |  |  |
| 13:1 ω1c | 0.6 | - | - | - | - | 2.2 | - | 18.0 |
| 14:1 ω5c | 0.1 | - | - | - | - | - | - |  |
| 15:1 ω5c | - | - | 0.6 | - | - | - | - | - |
| 15:1 ω8c | 0.2 |  |  | - | - | - | - | - |
| 17:1 ω8c | 3.0 | - | 0.9 | 0.9 | - | - | - | - |
| 18:1 ω9c | 32.5 | 21.4 | 17.5 | 22.8 | 19.3 | 20.3 | 16.6 | - |
| 20:1 ω9c | 0.3 | - | - | - | - | - | - | - |
| 20:4 ω6,9,12,15c | 1.7 | 3.3 | 3.4 | 2.8 | - | 2.6 | 3.7 | 1.9 |
| **Branched-chain** |  |  |  |  |  |  |  |  |
| 15:0 iso | 0.3 | - | - | - | - | - | - | - |
| 15:0 anteiso | 0.4 | - | 0.6 | - | - | - | - | - |
| 16:0 iso | 0.3 | - | - | - | - | - | - | - |
| 17:0 iso | 0.5 | - | 0.6 | - | - | - | - | - |
| 17:0 anteiso | 0.8 | 0.9 | 0.9 | 0.9 | - | - | - | 0.9 |
| 18:0 iso | 0.3 | - | - | - | - | - | - | - |
| 19:0 iso | 0.4 | - | - | - | - | - | - | - |

|  |  |  |  |  |  |  |  |  |  |
| --- | --- | --- | --- | --- | --- | --- | --- | --- | --- |
|  | **20:0 iso** | - | - | 0.4 | - | - | - | - | - |
|  | **Summed features\*** |  |  |  |  |  |  |  |  |
|  | **16:1 ω6c or 16:1 ω7c** | 1.4 | 2.1 | 2.2 | 1.8 | - | 1.8 | 2.6 | 1.7 |
|  | **18:1 ω6c or 18:1 ω7c** | 6.1 | 8.5 | 9.0 | 8.0 | 6.1 | 7.5 | 9.5 | 9.2 |
|  | **18:2 ω6,9c or 18:0 anteiso** | 5.9 | 10.1 | 11.1 | 9.5 | 8.7 | 8.0 | 11.2 | 9.9 |
|  | **19:1 ω9c or 19:1 ω11c** | 0.3 | - | - | - | - | - | - | - |

\*groups of more than one fatty acid that could not be separated by gas chromatography

For fatty acid analysis, cells of all strains were harvested from the same culture conditions during the late log phase (at 37 °C in a 5% CO2-enriched environment for 2 days on TSA with 5% sheep blood agar). Fatty acids were extracted from cells using the standard midi protocol (Sherlock Microbial Identification System, v6.0B), analysed with a gas chromatograph (6890 Series GC System, Hewlett Packard), and identified using the TSBA6 database of the Microbial Identification System [32]. The cellular fatty acid profiles of all strains included saturated, unsaturated, and branched-chain fatty acids (**Table 4**). The major fatty acids identified in *C. tuberculostearicum* ATCC 35692^T^, *C. accolens* ATCC 49725^T^, *C. hallux* sp. nov. CTNIH22^T^, and strains of *C. nasorum* sp. nov. and *C. yonathiae* were 18:1 ω9c (oleic acid), 16:0 (palmitic acid), and 18:0 (stearic acid). *C. hallux* sp. nov. CTNIH22^T^ had a higher amount of 18:1 ω9c (oleic acid) than other strains tested, and this fatty acid was absent from the composition of *C. macginleyi* ATCC 51787^T^. Several fatty acids were uniquely present in lesser amounts in *C. hallux* sp. nov. CTNIH22^T^ (e.g., 12:0, 15:1 ω8c, 20:1 ω9c).

### Description of *Corynebacterium hallux* sp. nov

*Corynebacterium hallux* sp. nov. (hal’lux. N.L. neut. n. *hallux* referring to the innermost toe, the skin site representing the source of this isolate).

Cells of *C. hallux* sp. nov. CTNIH22^T^ are Gram-positive, catalase-positive, oxidase-negative, non-spore-forming, non-motile bacilli (0.6-2.0 µm long and 0.4-0.6 µm wide). Optimal growth on solid medium was observed on BHI with 1% Tween 80 agar with aerobic incubation at 37 °C in a 5% CO_2_-enriched environment. Colonies on this medium were creamy white and measure approximately 3-5 mm in diameter; growth is weaker on TSA with 5% sheep blood agar, with non-hemolytic colonies measuring 1-2 mm in diameter. In liquid culture, *C. hallux* sp. nov. CTNIH22^T^ requires the addition of 0.2% Tween 80 for growth in BHI or TSB and tolerates salinity up to 14%. However, it has more stringent requirements for pH, with growth only observed at pH between 7.0 and 8.0. On biochemical testing, a positive reaction is observed for alkaline phosphatase with a weakly positive reaction for pyrrolidonyl arylamidase. No carbohydrate fermentation is noted in testing performed using API CORYNE strips. The major fatty acids identified are oleic (C18:1 ω9c; 32.5%), palmitic (C16:0; mean of 26.9%), and stearic (C18:0; 12.9%) acids. The genome size and DNA G+C content of the type strain are 2.53 Mb and 58.4 mol%, respectively.

The type strain, CTNIH22^T^ (=ATCC TSD-435^T^=DSM 117774^T^), was isolated from the toe web of a healthy adult. The partial 16S rRNA gene sequence of strain CTNIH22^T^ is available in GenBank (accession number: PQ252679). The GenBank accession number for the genomic sequence of this strain is GCF_032821755.1.

### Description of Corynebacterium nasorum sp. nov

*Corynebacterium nasorum* sp. nov. (nas’or.um L. gen. adj. *nasorum* referring to “of noses”, the human body site that is the source of the isolates).

Cells are Gram-positive, catalase-positive, oxidase-negative, non-spore-forming, non-motile bacilli (0.6-2.2 µm long and 0.4-0.6 µm wide). Optimal growth on solid medium is observed on BHI with 1% Tween 80 agar with aerobic incubation at 37 °C in a 5% CO_2_-enriched environment. Colonies on this medium are creamy white and measure 5-10 mm in diameter; growth is weaker on TSA with 5% sheep blood agar with non-hemolytic colonies. Optimal growth of *C. nasorum* sp. nov. in liquid medium is observed in BHI broth with 0.2% Tween 80; growth is also observed in TSB with 0.2% Tween 80. Growth of strains of *C. nasorum* sp. nov. occurs at pH between 6.0 and 8.0 and at salinity up to 10%. On biochemical testing, a positive reaction is observed for alkaline phosphatase, with a weakly positive reaction for pyrazinamidase and variable pyrrolidonyl arylamidase activity. No carbohydrate fermentation is noted in testing performed using API CORYNE strips. The major fatty acids are palmitic (C16:0; mean of 35.0%), oleic (C18:1 ω9c; mean of 19.5%), and stearic (C18:0; mean of 13.9%) acids. The genome size and DNA G+C content of the type strain are 2.46 Mb and 58.5 mol%, respectively.

The type strain, KPL3804^T^ (=ATCC TSD-439^T^=DSM 117767^T^), was isolated from a swab of the nostrils of a healthy adult aged between 31 and 60 years in Massachusetts, USA. The partial 16S rRNA gene sequence of strain KPL3804^T^ is available in GenBank (accession number: PQ149068). The GenBank accession numbers for the genomic sequences of the *C. nasorum* sp. nov. strains described in this study are GCF_037908315.1 (KPL3804^T^) and GCF_030229765.1 (MSK185).

### Description of strains of the recently described species *C. yonathiae*

Cells are Gram-positive, catalase-positive, oxidase-negative, non-spore-forming, non-motile bacilli (up to 5 µm in length and 0.4-0.6 µm wide). Optimal growth on solid medium is observed on BHI with 1% Tween 80 agar with incubation aerobically at 37 °C in a 5% CO_2_-enriched environment. Colonies on this medium are raised, creamy colonies between 5-10 mm in diameter; growth is weaker on TSA with 5% sheep blood agar, with non-hemolytic colonies measuring 2-3 mm in diameter. In liquid culture, strains require the addition of 0.2% Tween 80 for growth in BHI or TSB and tolerate salinity up to 10% and pH between 7.0 and 8.0. On biochemical testing, positive reactions are observed for alkaline phosphatase and pyrrolidonyl arylamidase. No carbohydrate fermentation is noted in testing performed using API CORYNE strips. The major fatty acids are palmitic (C16:0; mean of 37.6%), oleic (C18:1 ω9c; mean of 21.1%), and stearic (C18:0; mean of 14.5%) acids. The genome sizes of *C. yonathiae* strains MSK136 and KPL2619 are 2.47 Mb and 2.35 Mb, with DNA G+C content of 58.5 and 58.6 mol%, respectively.

## AUTHOR STATEMENTS

## 1.6 Authors and contributors

Writing – original draft: E.B.P., M.S.K., K.P.L., T.H.T.

Writing – review and editing: E.B.P., E.B., A.K.S., J.A.S., H.H.K., S.C., T.H.T., I.F.E., A.Q.R., K.P.L., M.S.K.

Investigation: E.B.P., T.H.T., I.F.E., E.B., A.K.S., N.A., A.Q.R., J.A.S., H.K., S.C., K.P.L., M.S.K.

Formal analysis: E.B.P., T.H.T., I.F.E., A.Q.R., M.S.K.

Resources: J.A.S., H.H.K., K.P.L., M.S.K.

Funding acquisition: M.S.K., K.P.L.

## 1.9 Ethical approval

The study protocol for collection of nasopharyngeal samples from infants in Botswana was approved by the Botswana Ministry of Health, the Princess Marina Hospital ethics committee, and institutional review boards at the University of Pennsylvania, Children’s Hospital of Philadelphia, McMaster University, and Duke University. Healthy volunteers were sampled as part of a prospective natural history study at the NIH Clinical Center approved by the NIH Institutional Review Board (www.clinicaltrials.gov/ct2/show/NCT00605878). The Forsyth Institutional Review Board approved the protocol (FIRB #17-02) used to collect nasal bacteria KPL strains in Massachusetts.

## 1.10 Consent for publication

Not applicable

## 1.11 Acknowledgements

We thank Dr. Wei Gao and Lukian Roberts for their efforts to isolate nasal *Corynebacterium* strains and prepare genomic DNA. This study utilized the computational resources of the NIH HPC Biowulf Cluster (http://hpc.nih.gov).

## ABBREVIATIONS

ANIb: average nucleotide identity based on BLAST+
ATCC: American Type Culture Collection
BHI: brain heart infusion
CO_2_: carbon dioxide
CNA: Colistin-Nalidixic Acid
FAME: fatty acid methyl ester
GTDB-Tk: Genome Taxonomy Database Toolkit
KEGG: Kyoto Encyclopedia of Genes and Genomes
MALDI-TOF MS: matrix-assisted laser desorption/ionization time-of-flight mass spectrometry
OD_600_: optimal density at a wavelength of 600 nanometers
PCR: polymerase chain reaction
rpm: revolutions per minute
rRNA: ribosomal RNA
TSA: tryptic soy agar
TSB: tryptic soy broth

## Supplementary Figures

**Figure S1.**
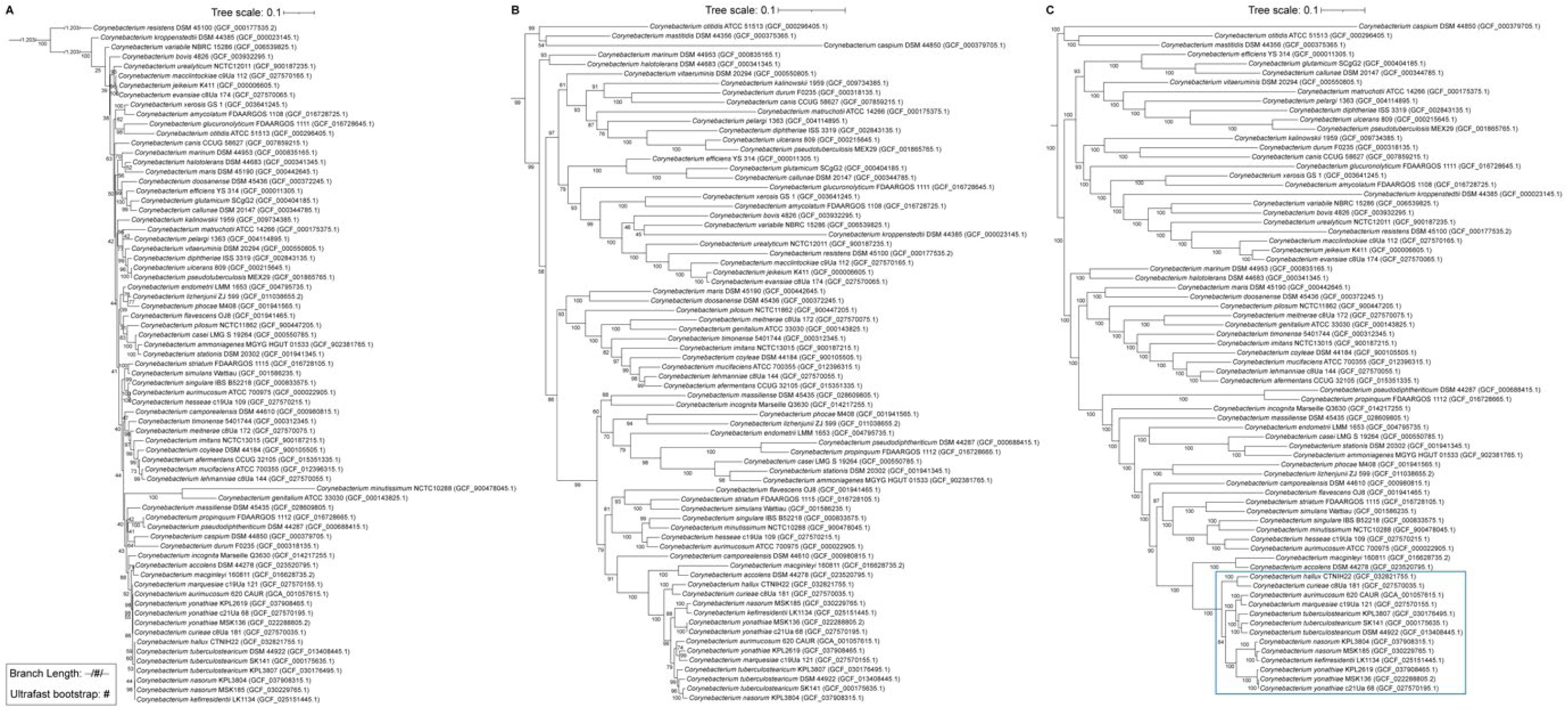
A phylogenomic tree of 68 members of the *Corynebacterium* genus illustrates the distinct species closely related to *Corynebacterium tuberculostearicum*. In contrast, phylogenetic trees of the same 68 *Corynebacterium species* based on either the (B) 16S rRNA gene or (C) *rpoB* gene showed the limitations of single-gene phylogenies for this genus. (**A**) A maximum-likelihood phylogenetic tree of 68 *Corynebacterium* species based on 16S rRNA gene sequences from 72 *Corynebacterium* strain genomes (**Table S1**) representing species across the breadth of the phylogeny of this genus has a number of poorly supported branches. (**B**) Although better than the full-length 16S rRNA gene phylogeny, a maximum-likelihood phylogeny based on full-length *rpoB* gene sequences of the same 72 strains also illustrates the limitation of single-gene phylogenies for resolving closely related *Corynebacterium* species. For example, the strains *C. yonathiae* KPL2619 (Cyo_KPL2619) and *C. marquesiae* c19Ua_121 have an average nucleotide identity based on BLAST+ (ANIb) below 95% (Figure 2), indicating they are distinct species, yet these incorrectly clade together here. Similarly, several strains assigned to the proposed species *C. nasorum* sp. nov. based on ANIb values are incorrectly in clades of other closely related species. (**C**) A maximum-likelihood phylogenomic tree of the same 68 *Corynebacterium* species was constructed using 193 concatenated and aligned shared single-copy core gene clusters from 72 *Corynebacterium* strain genomes (**Table S1**). The majority of branches in this phylogeny exhibit strong support with ultrafast bootstrap values of 95 or higher. See Supplemental Methods for description of the construction of these phylogenies.

**Figure S2.**
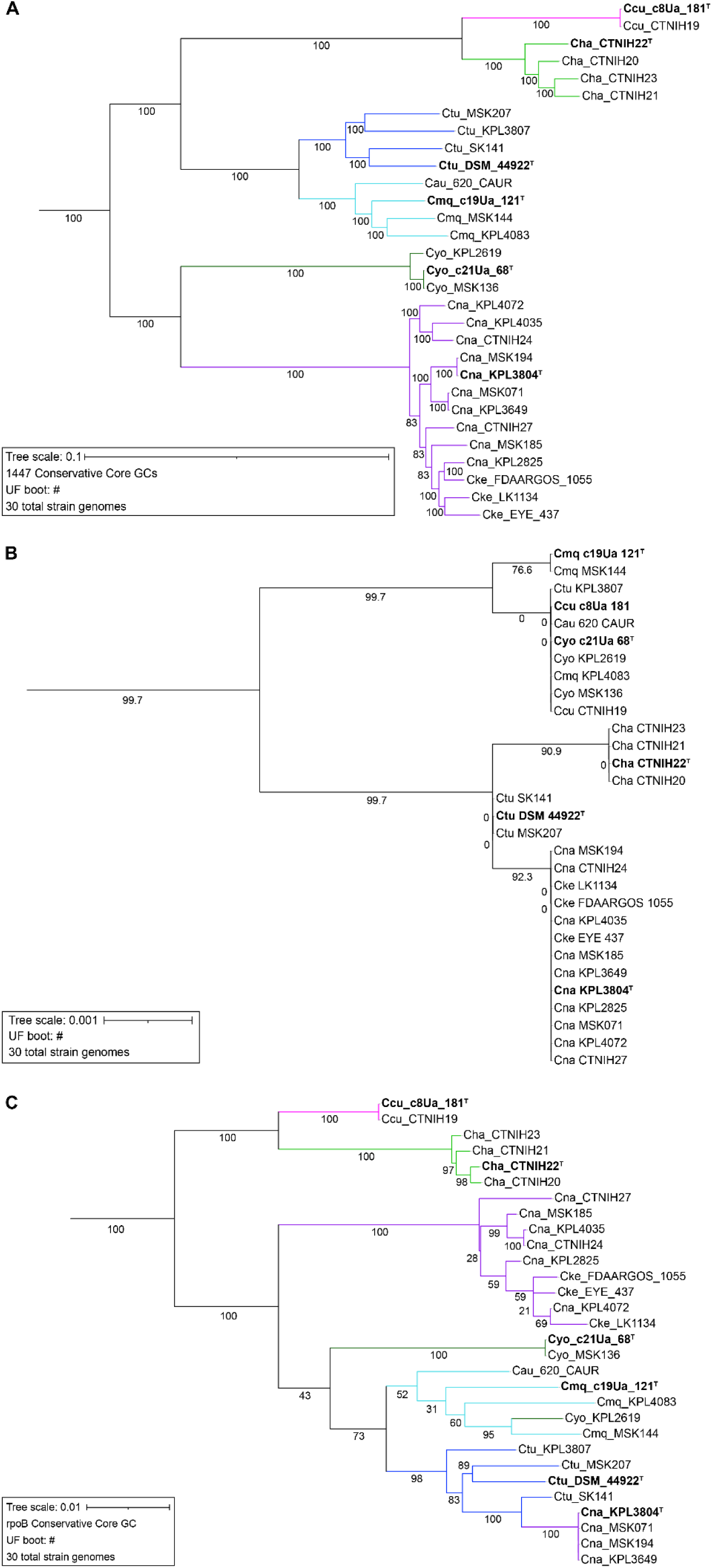
A phylogenomic tree provided superior resolution among the species most closely related to *Corynebacterium tuberculostearicum* compared to single-gene phylogenies. **(A)** The maximum likelihood phylogenomic tree constructed based on 1447 conservative core gene clusters provided higher resolution of the distinct clades within the *Corynebacterium tuberculostearicum* species complex. The use of a large pool of single copy gene clusters shared across 30 *Corynebacterium* genomes within the *C. tuberculostearicum* species complex enhanced species delineation, with robust ultrafast bootstrap values supporting the distinct clades. Ccu (pink branches) is *C. curieae*; Cha (light green branches) is *C. hallux* sp. nov.; Ctu (blue branches) is *C. tuberculostearicum*; Cmq (turquoise branches) is *C. marquesiae*; Cyo (dark green branches) is *C. yonathiae*; and Cna (purple branches) is *C. nasorum* sp. nov. **(B)** In contrast, a full-length 16S rRNA gene maximum likelihood phylogeny had strains of different species sometimes intermingled in a single clade and was poorly supported based on ultrafast bootstrap values. This is consistent with the known limitations of using 16S rRNA gene phylogenies within this genus and highlights the limitations of using the 16S rRNA gene for resolving evolutionary relationships within the *C. tuberculostearicum* species complex. **(C)** The full-length *rpoB* gene maximum likelihood phylogeny had better support than the 16S rRNA gene for the species within the *C. tuberculostearicum* species complex. However, it still had two clades with intermingled species and was inferior to the phylogenomic tree using conservative core gene clusters shown in S2A.

## Supplementary Methods

### Construction of phylogenic trees

To generate the maximum likelihood 16S rRNA gene phylogenies shown in Figures 1, S1A, and S2B, we performed the following steps. First, to identify the 16S rRNA genes present in each genome, we ran barrnap v0.9 (https://github.com/tseemann/barrnap) with default parameters on the fasta files containing the genome assemblies for each phylogeny. We then used seqkit (v2.6.0) grep -r -n -p ‘16S_rRNA’ to select the 16S rRNA gene sequences from each genome’s total rRNA sequences [33]. For genomes that had multiple copies of the 16S rRNA gene, we manually inspected the sequences and removed copies that were less than 50% of the expected 16S rRNA gene sequence length using AliView v1.28 [34], aligned the remaining copies using MUSCLE v3.8.1551 with default parameters, and generated a consensus 16S rRNA gene sequence using the EMBOSS cons command [35, 36]. We concatenated and aligned the single and consensus 16S sequences with the linux ‘cat’ command and MUSCLE. The resulting 16S rRNA gene alignment was used as input for IQ-TREE2 v2.1.3 [37] and we set the parameters -alrt to 1000 and -B to 1000.

To generate the maximum-likelihood *rpoB* phylogenies shown in Figures S1B and S2C, we identified the single copy *rpoB* gene cluster from the conservative core determined with GET_HOMOLOGUES [38] (see below) and then aligned and concatenated the *rpoB* gene from all the genomes with GET_PHYLOMARKERS [39]. Then we used IQ-TREE2 with the same parameters as the 16S rRNA tree [37].

To generate the maximum likelihood phylogenomic trees in Figures 3, S1C and S2A, we used Prokka v1.14.6 [40] with default settings to annotate each bacterial genome, based on the prediction of coding sequences with Prodigal [41]. For detailed methods on the annotation of genomic assemblies, please see https://klemonlab.github.io/NovCor_Manuscript/Methods_Prokka_Annotations.html. We then used GET_HOMOLOGUES (version 13062023) [38] to separately identify the core gene clusters (GCs) shared by the set of strains used for each individual tree. The consensus of the single copy core GCs from three clustering algorithms; bidirectional best-hits, cluster of orthologs triangles (COGS) v2.1 [42], and Markov Cluster Algorithm OrthoMCL (OMCL) v2.4 [43], defined the conservative shared core genome for each group using ./get_homologues.pl. Subsequently, we employed GET_PHYLOMARKERS v2.2.9.1 [39] to align and concatenate the shared single copy core gene clusters. These were then analyzed using IQ-TREE2 v2.1.3 [37] with the following parameters: -p (edge-linked partition model and ModelFinder functions) [44, 45], -alrt 1000 (replicate SH-like approximate likelihood ratio test) [46], and -B 1000 (number of ultrafast bootstrap replicates) [21].

To visualize, scale, edit, annotate names, and root trees at the midpoint for each phylogeny, we used the phylogenetic tool iTOL version 6 [47]. For detailed methods on the construction of all phylogenetic trees, please see https://klemonlab.github.io/NovCor_Manuscript/Methods_Phylogenies.html.

### Average Nucleotide Identity

We used pyani (version 0.2.9) [22] with ANIb BLAST+ [23, 48] to construct a WGS identity ANI heat matrix. We used the data in the pyani output file ANIb_percentage_identity.tab to create Figure 2 in R, with genome order based on the corresponding .svg file. For detailed methods, see https://klemonlab.github.io/NovCor_Manuscript/Methods_ANIs.html.

